# ORBIT: Orthogonal Rotation for Biological Inter-species Transfer

**DOI:** 10.64898/2026.05.04.722193

**Authors:** Paul Wissenberg, Jia Min Lee, Marek Mutwil

## Abstract

**Motivation:** Cross-species gene embeddings are central to transferring functional annotations between species. A recent method demonstrated that species-specific STRING (PPI) network embeddings can be aligned across 1322 eukaryotes with autoencoders (FedCoder), but this approach is computationally expensive, depends on careful hyperparameter selection, leaves substantial room for improvement in cross-species retrieval quality, and has not been demonstrated on coexpression networks.

**Results:** We introduce an alignment pipeline for cross-species coexpression network embeddings based on orthogonal Procrustes rotation. Species-specific Node2Vec embeddings of coexpression networks are aligned to a shared space using ortholog anchors from OrthoFinder, solved in closed form via Singular Value Decomposition (SVD). Applied to 153 plant species and 5.7 million genes, Procrustes alignment achieves four-fold higher cross-species Spearman correlation and consistently higher retrieval metrics than the SPACE autoencoder, while leaving within-species coexpression structure invariant (preservation ratio 1.000 against the unaligned baseline). The full alignment completes in under three minutes on a single CPU, and on downstream tasks, Procrustes embeddings improve within-species GO term prediction and outperform SPACE for cross-species GO transfer. Procrustes and sequence embeddings remain complementary for biological-process prediction, consistent with observations from SPACE.

**Availability:** Code for producing the embeddings is made available at https://github.com/pwissenberg/orbit.

## 1. Introduction

*Arabidopsis thaliana*, the model flowering plant, has 18,742 manually curated UniProtKB/Swiss-Prot entries ((UniProt Consortium 2023), accessed April 2026); non-model species such as *Picea abies* (Norway spruce) have just 30, a 625-fold annotation gap. This asymmetry constrains the entire field of plant functional genomics: due to often many-to-many relationships between plant orthologs, experimental knowledge accumulated in *A. thaliana* cannot be straightforwardly transferred to crops or wild relatives, where new annotations are most urgently needed for crop improvement and stress biology. The Critical Assessment of Function Annotation (CAFA), held triennially, has formalized this as a community benchmark for predicting protein function from sequence and structure (Friedberg and Radivojac 2017). Yet sequence similarity alone misses much of the regulatory and pathway context that distinguishes paralogs and homologs, motivating the use of gene network embeddings as a complementary substrate for cross-species function transfer.

Protein language models such as ProtT5 (Elnaggar *et al*. 2022), trained on tens of millions of protein sequences, encode each protein as a fixed-dimensional vector that captures both sequence and structural context. Because these models embed sequences from any species into the same vector space, they are inherently cross-species comparable and have become a standard input for tasks such as subcellular localization (Thumuluri *et al*. 2022) and protein function prediction (Yao *et al*. 2021). Network-derived embeddings provide complementary information: graph representation learning methods such as Node2Vec (Grover and Leskovec 2016) embed each gene by its connectivity in a coexpression or protein-protein interaction network, encoding the functional and regulatory neighborhood that sequence alone cannot reveal. Unlike sequence embeddings, however, network embeddings are learned independently for each species, so the same biological function maps to arbitrary coordinates in different organisms, making cross-species comparison impossible without an explicit alignment step.

Aligning network embeddings across species is a non-trivial representation learning problem. The goal is to find a transformation that places orthologous genes close together while preserving the within-species structure of each network. Early work used closed-form alignment of kernel-based embeddings to map species-specific networks into a shared space via homolog landmarks (Fan *et al*. 2019). More recent methods learn a mapping into a shared latent space using autoencoders (Li *et al*. 2023), most notably SPACE (Hu *et al*. 2025), which uses the FedCoder framework to align approximately 1,322 eukaryotic species embedded from STRING. An alternative is to learn a single joint embedding across species using ortholog pairs as edge constraints during training (Mancuso *et al*. 2024). Among these, SPACE is the current community reference. However, FedCoder learns its mapping by gradient descent over an unconstrained linear layer, which is computationally expensive and does not by itself preserve within-species distances.

We make four contributions to this work. First, we present a training-free alignment pipeline for coexpression-based gene embeddings spanning 153 plant species, anchored by Jaccard-weighted ortholog pairs. Second, in a direct comparison against SPACE on the same species, we demonstrate a four-fold improvement in cross-species Spearman correlation (Wilcoxon p < 10^−2^□), a 500-fold reduction in runtime while using a CPU only. Third, we evaluate the resulting embeddings on within-species and cross-species GO term prediction and on subcellular localization, clarifying when coexpression embeddings provide signal beyond protein language models. Fourth, we release pre-aligned embeddings as a public resource for the plant functional genomics community.

## 2. Methods

### 2.1 Data

For the coexpression networks of 153 plant species, we use the TEA-GCN method (Lim *et al*. 2025), selecting the top 50 edges per gene ranked by mutual rank z-score. The orthologs and orthogroups were identified with the OrthoFinder v3.1.0 (Emms and Kelly 2019). Protein sequence embeddings were generated with ProtT5-XL-UniRef50 (Elnaggar et al. 2022), averaged per protein to obtain a single embedding per protein, following Hu et al. (2025) for direct comparability. Per-species counts of manually curated proteins reported in the Introduction were retrieved from UniProtKB/Swiss-Prot (UniProt Consortium 2023) via the EBI Proteins API using the reviewed=true&taxid=<NCBI_taxid> filter on 25 April 2026. For the seed species, we selected five phylogenetically diverse species spanning major plant lineages, arranged from most recent to oldest divergence: *Arabidopsis thaliana* (eudicot), *Oryza sativa* (monocot), *Picea abies* (gymnosperm), *Selaginella moellendorffii* (lycophyte), and *Marchantia polymorpha* (liverwort).

### 2.2 Species-specific embeddings (Node2Vec)

We generated 128-dimensional species-specific embeddings using weighted Node2Vec via PecanPy (Liu and Krishnan 2021), following Hu et al. (2025). Hyperparameters were selected via grid search, optimizing cross-species Spearman correlation across seed species pairs (Table S1, Fig. S1). The best configuration (p = 1.0, q = 0.7) was robust across parameter ranges (ρ = 0.502– 0.526).

### 2.3 Cross-species alignment by Procrustes rotation

Each species’ Node2Vec embedding represents its genes in an independent 128-dimensional space. Because these spaces are learned separately, identical network neighborhoods are likely to map to arbitrary coordinates in different species, making direct comparison impossible. However, orthologous genes identified by OrthoFinder provide known correspondences between species. We exploit these anchor pairs to find an optimal rotation that maps one species’ embedding space onto another, placing orthologs close together while preserving each species’ internal structure. We apply orthogonal Procrustes rotation, previously used to align learned word embeddings across languages (Artetxe, Labaka, and Agirre 2016), to align gene coexpression embeddings across species.

Formally, given two species A and B with embedding matrices *X*_*A*_ ∈ *ℝ*^*n × d*^ and *X*_*B*_ ∈ *ℝ*^*m × d*^, and *k* ortholog pairs forming anchor matrices _*A B*_ ∈ *ℝ*^*k × d*^, we seek the orthogonal matrix R that minimizes the distance between aligned orthologs:

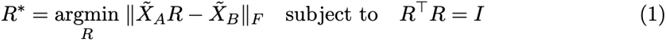

This has a closed-form solution via singular value decomposition:

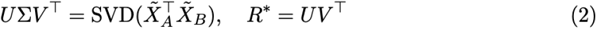

The rotation is then applied to all genes in species A to project them into the shared space:

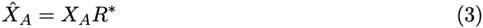

Because R is orthogonal, all pairwise distances and cosine similarities within a species are preserved exactly.

To extend this pairwise alignment to all 153 species, we first align all five seed species to the main model plant *A. thaliana* as the reference, chosen for its superior functional annotation. Each seed species’ rotation matrix is computed independently against *A. thaliana*, placing all seeds in a shared space. Each of the remaining 148 non-seed species is then aligned to its nearest seed, determined by orthogroup overlap. Because all seeds are already aligned to the same reference, transitivity places all species in a single shared embedding space.

Orthogroups often contain multiple genes per species, producing many-to-many ortholog mappings. To prioritize the most informative anchor pairs, we weight each ortholog pair (g_A_, g_B_) by the Jaccard similarity of their orthogroup-mapped coexpression neighborhoods N_O_(g_A_) and N_O_(g_B_)

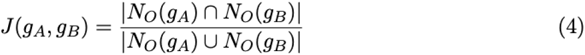

where N_O_(g) denotes the set of orthogroups represented among the coexpression neighbors of gene g. Mapping to orthogroups makes the neighborhoods comparable across species. B using Jaccard similarity, pairs with highly conserved coexpression context receive more influence on the rotation.

Once the rotation R has been computed and applied, retrieval of cross-species ortholog candidates uses cosine similarity in the aligned space. To mitigate hubness — the phenomenon where some genes become nearest neighbors of many others in high-dimensional spaces — we apply cross-domain similarity local scaling (CSLS; Conneau *et al*. 2018) during retrieval. For a query gene g_A_ in species A and a candidate gene g_B_ in species B:

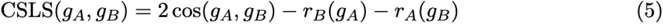

where r_B_(g_A_) is the mean cosine similarity of g_A_ to its K nearest neighbors in species B (with neighborhoods N (gA) defined by cosine similarity in the aligned space):

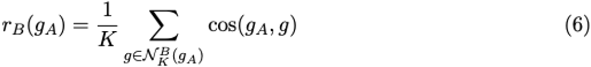

and r_A_(g_B_) is defined symmetrically. This penalizes genes that act as hubs in either species and rewards mutual nearest neighbors. We use K=10 following Conneau *et al*. (2018), applied to L2-normalized embeddings, both during iterative anchor refinement and final cross-species retrieval.

We chose orthogonal rotation over a learned mapping for three reasons: it preserves all within-species distances, it requires no training or hyperparameter tuning, and the rotation is obtained in closed form by SVD, replacing iterative gradient-descent training with a single non-iterative step. This assumes that the optimal cross-species mapping is well-approximated by an orthogonal transformation. This assumption is well-established for cross-lingual word embeddings (Artetxe, Labaka, and Agirre 2016) and, as we show in Section 3.2, holds equally for plant gene coexpression embeddings.

### 2.4 SPACE autoencoder baseline (comparison method)

For a like-for-like comparison, we retrained the SPACE autoencoder (Hu *et al*. 2025) on the same 153-plant Node2Vec embeddings used for Procrustes alignment, using the official FedCoder implementation with the published default hyperparameters: 512-dimensional shared latent space, alpha = 0.5 reconstruction/alignment weighting, and 500 epochs. We deliberately did not re-tune SPACE hyperparameters — using the authors’ published settings provides a principled baseline and avoids unfair degradation against a competing method. Throughout this paper, “SPACE” refers to this retrained 153-plant model unless stated otherwise.

### 2.5 Projecting non-seed species into shared embedding space

After aligning all five seed species by rotating each onto *A. thaliana*, we project the remaining 148 non-seed species into the shared space. Each non-seed is aligned to its nearest seed, determined by Jaccard similarity given in (4). This ensures more shared anchors are available for computing the rotation. Because each projection is independent, this step parallelizes trivially across all 148 non-seed species. Because all seeds are already in the same space, transitivity ensures a single shared embedding for all 153 species.

### 2.6 Alignment quality metrics

To evaluate alignment quality, we follow the SPACE evaluation framework for direct comparability. Cross-species retrieval is assessed by ranking all genes in species B by cosine similarity to a query gene in species A and measuring whether known orthologs are retrieved, using Hits@50 (the fraction of query genes whose best Jaccard-weighted ortholog partner appears among the top 50 nearest neighbors in the target species), MRR@50 (the mean reciprocal rank of that best Jaccard-weighted partner, where rank r ≤ 50 contributes 1/r and absence contributes 0), Top-M Hits@50 (where M = 10 corresponds to the 10 highest-Jaccard ortholog partners; the criterion is whether any of these appears in the top 50 retrieved), and Spearman ρ between cosine similarity and Jaccard weight, computed over ortholog pairs. Ground truth ortholog pairs are weighted by the Jaccard similarity of their coexpression neighborhoods, mapped to orthogroups. Within-species preservation is measured by Precision@50 and Recall@50 on coexpression network reconstruction. To test statistical significance, we apply Mann-Whitney U tests on cosine similarity distributions of orthologous versus non-orthologous pairs, and Wilcoxon signed-rank tests on paired metrics across species.

### 2.7 Downstream tasks

To evaluate the biological utility of aligned embeddings, we performed three downstream prediction tasks. For within-species GO prediction, we trained logistic regression classifiers on *A. thaliana* using the NetGO 2.0 benchmark (∼13k proteins), reporting Fmax across molecular function, biological process, and cellular component (Yao *et al*. 2021). For subcellular localization, we used the DeepLoc 2.0 benchmark (∼4,500 proteins), reporting F1-micro (Thumuluri *et al*. 2022). For cross-species GO transfer, we trained on *A. thaliana* GO annotations filtered to experimental evidence codes from UniProt GOA (UniProt Consortium 2023) and predicted gene functions in rice, maize, soybean, and Medicago. All tasks compared five embedding configurations: SPACE, Procrustes, ProtT5, SPACE+ProtT5, and Procrustes+ProtT5.

### 2.8 Implementation and availability

The alignment pipeline is implemented in Python, using NumPy and SciPy for SVD computation and PecanPy for Node2Vec embedding generation (Liu and Krishnan 2021). All experiments were run on a single machine with 48 CPU cores and 4× NVIDIA L4 GPUs (23 GB each). Procrustes alignment used CPU only; the SPACE autoencoder used GPU. All code and the pre-aligned embeddings for 5 species are available under https://github.com/pwissenberg/orbit.

## 3. Results

### 3.1 Procrustes rotation aligns plant coexpression embeddings across 153 species

In this paper, we demonstrate that a simple rotation outperforms a trained autoencoder for aligning cross-species coexpression embeddings. We computed coexpression embeddings for 153 plant species and 5.7 million genes, generated with orthogonal Procrustes rotation and the SPACE autoencoder, as well as ProtT5 sequence embeddings. Our approach leverages coexpression networks from TEA-GCN and orthology relations from OrthoFinder to create cross-species gene embeddings. The workflow begins by generating 128-dimensional species-specific embeddings using Node2Vec on each coexpression network (Fig. 1). These embeddings are then aligned across species using a two-step process. First, five phylogenetically diverse seed species are mutually aligned using orthogonal Procrustes rotation. This process computes optimal rotation matrices from ortholog pairs via SVD, mapping each seed species into a shared embedding space while preserving within-species coexpression structure exactly. Subsequently, each of the remaining 148 non-seed species is aligned to its nearest seed using the same rotation procedure. We evaluated biological pathway integrity with KEGG pathway co-membership ROC curves (Kanehisa *et al*. 2021), which revealed that the alignment did not degrade pathway-level signal (Fig. S2).

**Figure 1.**
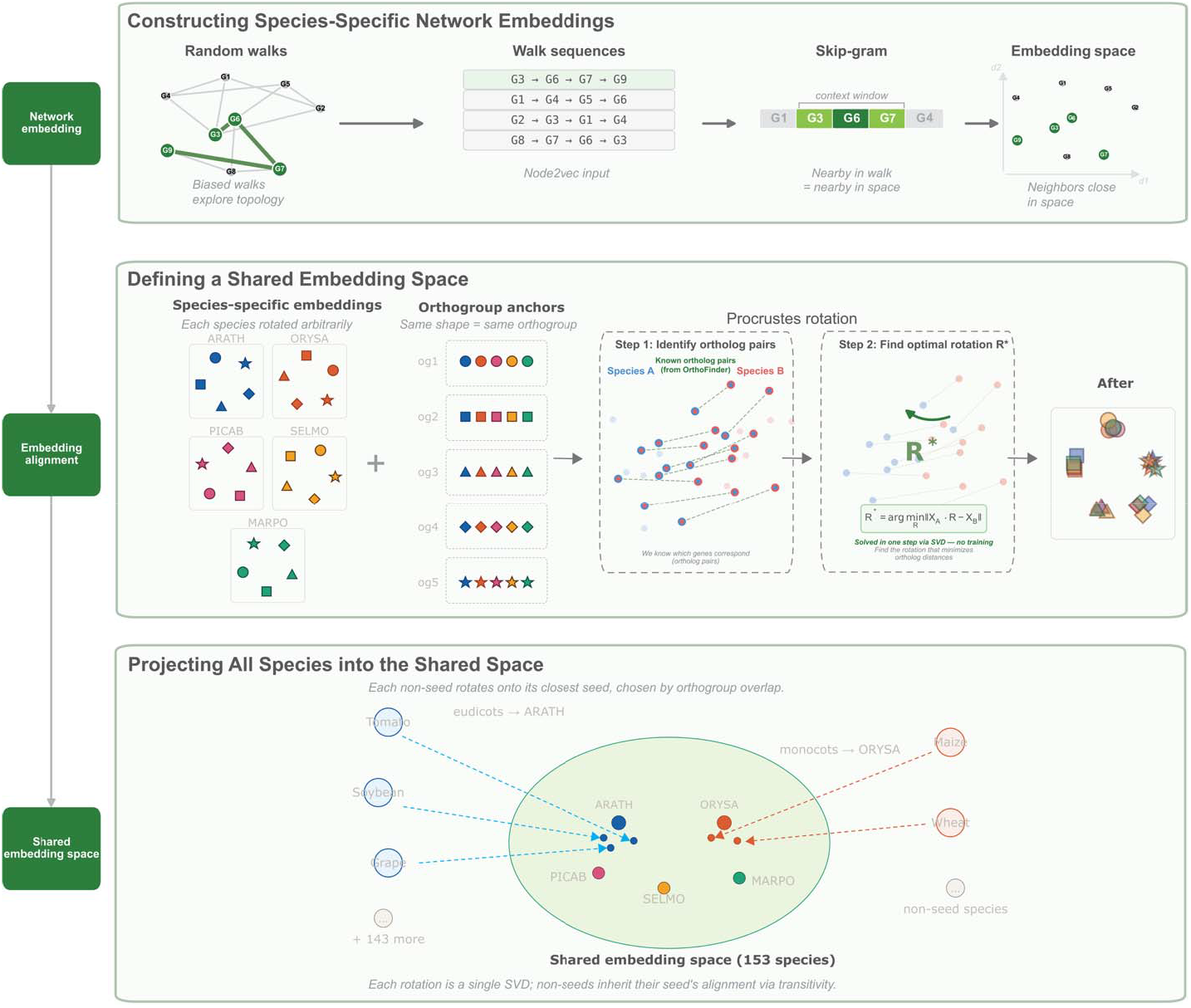
Pipeline for cross-species gene function transfer using coexpression network alignment. Node2Vec generates species-specific gene embeddings by learning from biased random walks on each coexpression network. Orthologous gene pairs identified by OrthoFinder anchor the cross-species alignment, mapping five seed species into a shared embedding space. All remaining species are projected into the shared space by aligning each to its nearest seed. Procrustes rotation uses ortholog pairs from OrthoFinder to define known correspondences between two species’ embeddings. The optimal rotation R* is found in closed form via SVD by minimizing the squared distance between paired orthologs across species. Applying R* rotates species A into species B’s space, placing orthologs close together while preserving each species’ within-species geometry by construction (rotation is an isometry).

### 3.2 Procrustes rotation outperforms SPACE autoencoder on cross-species retrieval

An effective cross-species alignment should place genes with conserved network context close together in the shared embedding space, irrespective of species of origin. To evaluate alignment quality, we compared Procrustes rotation against the SPACE autoencoder on cross-species retrieval metrics across 153 species pairs (Fig. 2A, Table S2). Procrustes achieves a Spearman correlation of 0.41 compared to 0.09 for SPACE, a four-fold improvement. Retrieval accuracy follows the same pattern (Hits@50: 0.13 vs 0.01; Top-M Hits@50: 0.19 vs 0.02). Within-species coexpression structure is preserved on par with the retrained SPACE-v2 baseline (Precision@50: 0.54 vs 0.55; Recall@50: 0.55 vs 0.55), confirming that the rotation does not distort local network neighborhoods. This parity is consistent with the theoretical guarantee that, by construction, orthogonal rotation leaves within-species geometry unchanged. The cross-species improvement is highly significant: a Wilcoxon signed-rank test on Spearman ρ across all 153 species pairs yields p = 5.6 × 10^−2^□, with Procrustes outperforming SPACE on every single pair (153/153). A component ablation isolating the contributions of Jaccard-weighted anchors, iterative refinement, and CSLS retrieval over basic orthogonal Procrustes is shown in Fig. S3. Alignment quality is essentially insensitive to ortholog anchor count — subsampling to 10% reduces mean Spearman ρ by only 1.1% (Fig. S4).

**Figure 2.**
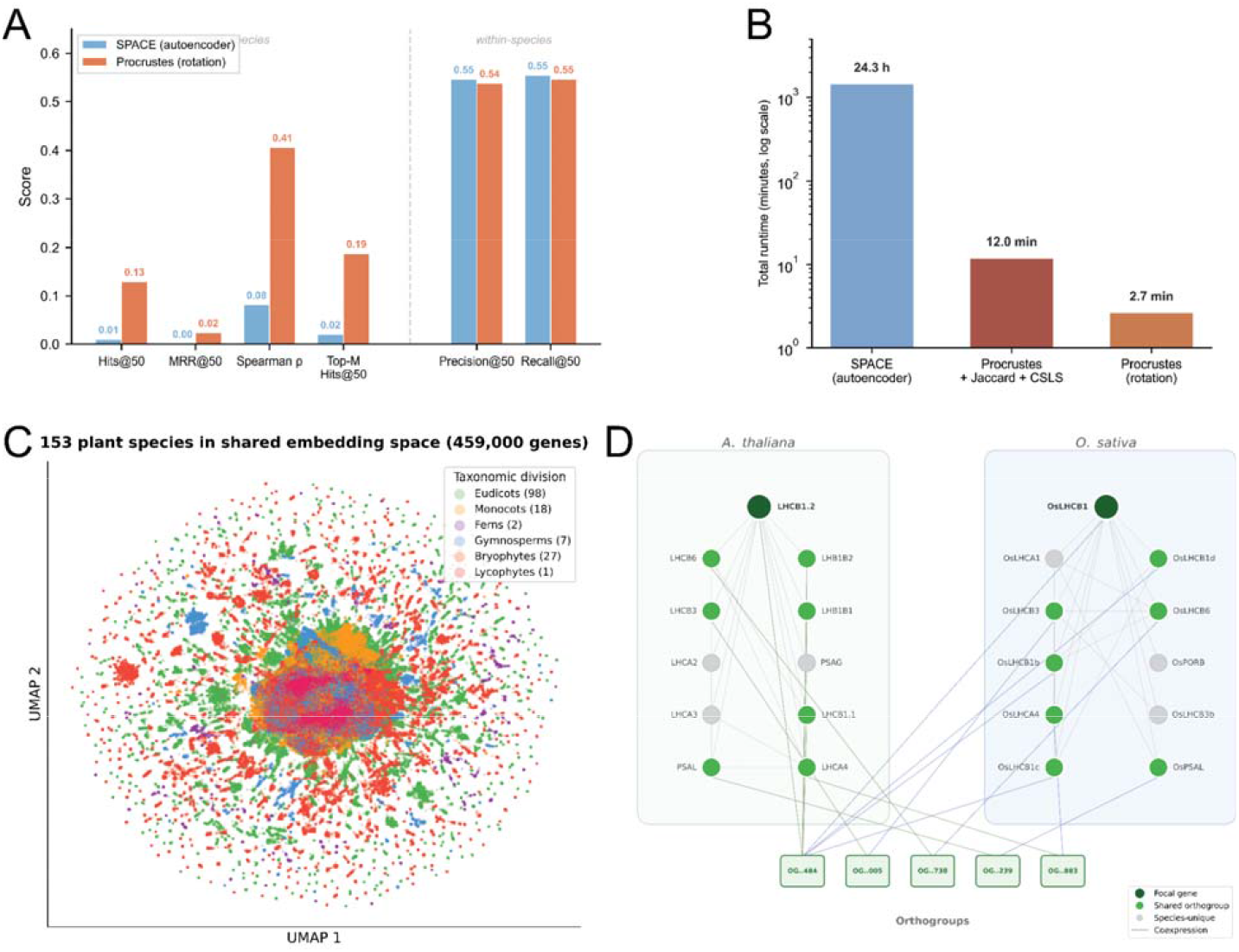
Alignment quality, runtime, and shared embedding space. (A) Cross-species retrieval metrics for Procrustes rotation and SPACE autoencoder across seed species pairs. (B) Runtime comparison (log scale). (C) UMAP projection of 459,000 genes (a subsample) from 153 species, colored by taxonomic division. (D) Conserved coexpression neighborhoods of LHCB1.2 (A. thaliana) and OsLHCB1 (O. sativa). Green: shared orthogroups; gray: species-unique. Bottom: shared orthogroups connecting the two networks.

### 3.3 Alignment completes in minutes on CPU

Beyond alignment quality, Procrustes rotation offers a substantial computational advantage (Fig. 2B, Table S3). The full alignment of 153 species completes in 2.7 minutes on a single CPU, compared to 24.3 hours for the SPACE autoencoder on GPU, which is more than a 500-fold speedup. Because SVD is constant-time in the number of species, adding new species to the shared space is near-instant as new networks become available.

### 3.4 The shared space recovers plant phylogeny

To visualize the shared embedding space, we projected 459,000 genes (a random subsample) onto two dimensions using UMAP (Fig. 2C). Genes cluster by taxonomic division, with eudicots and monocots forming the central mass and earlier-diverging lineages such as bryophytes and lycophytes positioned at the periphery. This confirms that the alignment preserves biologically meaningful structure rather than collapsing species into a single point cloud. The biological basis for this alignment is illustrated by the conserved coexpression neighborhoods of LHCB1.2 and OsLHCB1, where 7 of 11 neighbors share orthogroups across species (Jaccard = 0.40), and the two focal genes achieve cosine similarity 0.92 in the aligned space (Fig. 2D).

### 3.5 Within-species GO prediction is improved in *A. thaliana*

We evaluated the utility of aligned embeddings for within-species gene function prediction on *A. thaliana* using the NetGO 2.0 benchmark (Fig. 3A). Procrustes’ embeddings outperform SPACE on cellular components (Fmax: 0.42 vs 0.37) and biological process prediction (0.14 vs 0.12), with molecular function comparable (0.78 vs 0.79). Concatenating ProtT5 sequence embeddings yields only marginal improvement (MF: 0.79 vs 0.78), suggesting that annotation coverage, rather than embedding capacity, limits performance.

**Figure 3.**
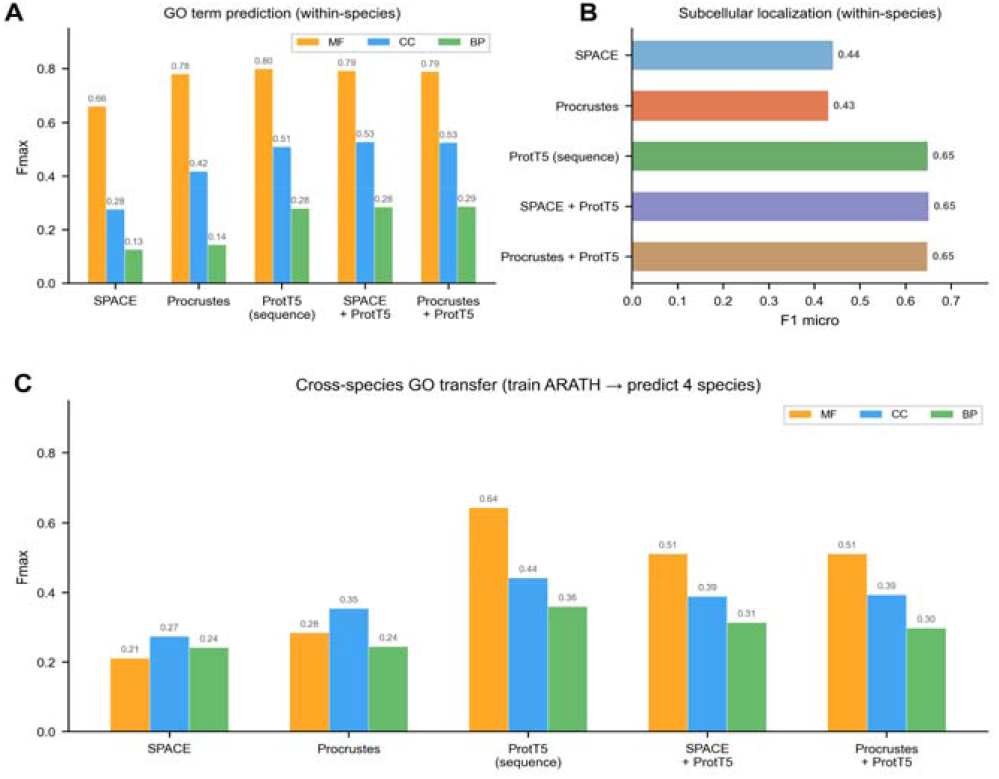
Downstream evaluation of aligned embeddings. (A) Within-species GO term prediction on A. thaliana (NetGO benchmark). (B) Subcellular localization prediction (DeepLoc 2.0 benchmark). (C) Cross-species GO term transfer from A. thaliana to rice, maize, soybean, and Medicago.

### 3.6 Subcellular localization is a sequence-dominated task

For subcellular localization prediction using the DeepLoc 2.0 benchmark, coexpression-based embeddings from Procrustes and SPACE achieve F1-micro of 0.45 and 0.47 respectively, while ProtT5 alone reaches 0.65 (Fig. 3B, Table S4; per-localization MCC in Fig. S5). Combining coexpression and sequence embeddings does not improve over ProtT5 alone. This is expected, as subcellular targeting is dominated by local sequence motifs rather than in coexpression relationships.

### 3.7 Procrustes embeddings improve cross-species GO transfer

To evaluate cross-species function transfer, the primary use case for aligned embeddings, we trained classifiers on *A. thaliana* GO annotations and predicted gene functions in rice, maize, soybean, and Medicago (Fig. 3C, Table S5; per-species breakdown by GO aspect in Fig. S6). Procrustes outperforms SPACE across all three GO aspects (MF: 0.28 vs 0.21; CC: 0.35 vs 0.27; BP: 0.24 vs 0.24). ProtT5 alone remains strongest for molecular function (0.64), where sequence largely determines function. However, for biological process and cellular component prediction, coexpression alignment contributes value that sequence alone cannot provide, and Procrustes delivers more of that value than SPACE.

## 4. Discussion

### Why rotation works so well

Both Procrustes and the SPACE FedCoder autoencoder learn linear transformations between species-specific embedding spaces; inspection of the FedCoder source code confirms a single linear layer without activation functions, consistent with the architectural choices reported in the SPACE paper. The difference lies in the constraint: Procrustes restricts the mapping to the orthogonal subset, for which the optimal solution is known analytically via singular value decomposition, while FedCoder optimizes over the full unconstrained linear space by gradient descent. When the optimal answer is known to lie in a constrained subset, gradient descent over the larger space is wasted effort. Empirically, this manifests as a four-fold improvement in cross-species Spearman correlation and exact preservation of within-species structure: the precision and recall at 50 of Procrustes-aligned embeddings match the raw Node2Vec embeddings to four decimal places (preservation ratio 1.0000), confirming the theoretical isometry guarantee.

### When autoencoders might still win

If embedding spaces are highly non-isometric, for example, due to very different network densities or asymmetric orthology distributions, an unconstrained linear or nonlinear mapping could, in principle, recover structure that orthogonal rotation cannot. Our results suggest this is not the regime for coexpression networks within Archaeplastida; the assumption that aligned orthologs differ by rotation alone holds well across the roughly 700-million-year span of plants. The picture may differ across kingdoms, where divergent network topologies and weaker ortholog signals could push the problem outside the orthogonal regime.

### Limitations

Several caveats apply. First, our scope is restricted to plants; the original SPACE study (Hu et al. 2025) covered ∼1,322 eukaryotes using STRING-derived networks. Before equivalent generality claims can be made, Procrustes alignment must be tested at that broader taxonomic scope and on PPI (rather than coexpression) data. Second, coexpression network coverage is uneven across the 153 species, with sparser networks in non-seed taxa limiting embedding quality regardless of alignment method. Third, ortholog anchor quality depends entirely on OrthoFinder, and systematic errors in orthogroup inference propagate into the rotation matrices. Fourth, our pipeline assumes a single global rotation per species pair; if different regions of the embedding space require different transformations, for example, to accommodate lineage-specific gene-family expansions, a single *R\** underfits the problem. Fifth, cross-species functional benchmarks are scarce: only *Oryza sativa* provides more than 500 test proteins with experimental evidence codes, limiting the statistical power of our downstream evaluation. We also note that, against the retrained SPACE-v2 baseline rather than vanilla SPACE, within-species structure preservation is similar between the two methods; the Procrustes advantage is concentrated in cross-species correspondence, where it dominates on 153 of 153 species pairs.

### Relationship to SPACE

This work is a methodological refinement with a narrower scope, not a replacement. SPACE remains the authoritative cross-kingdom resource based on STRING and represents an important contribution to comparative functional genomics across all eukaryotes. Our results show that, within plants, and on coexpression rather than PPI data, a far simpler method is both faster and more accurate, but the broader architectural decision in SPACE to provide a unified embedding for any eukaryotic protein remains valuable for use cases that span kingdoms.

### Broader implications

Closed-form alignment removes the principal computational barrier to maintaining a continuously updated cross-species embedding space. As new biological networks are released, new species can be incorporated in minutes via a single SVD, rather than requiring a GPU-intensive training. This has immediate value for annotation transfer to previously inaccessible non-model lineages. More broadly, our results illustrate a methodological principle worth flagging in an era of expressive learned models: when the optimal hypothesis class is known, restricting to that class often beats searching the larger space, both in solution quality and in computational cost.

## 5. Conclusions

Peter Schönemann published the closed-form solution to the orthogonal Procrustes problem in 1966, building on a least-squares construction Hurley and Cattell had described twenty years earlier. Wahba reused it for satellite attitude estimation. Kabsch reused it for protein structure alignment. Now it is our turn to use it for cross-species coexpression network alignment.

We have shown that closed-form orthogonal Procrustes rotation is sufficient to align coexpression-network embeddings across 153 plant species, outperforming the SPACE autoencoder by four-fold on cross-species retrieval and by more than 500-fold in runtime while preserving within-species structure exactly. This improvement carries through to downstream tasks: Procrustes embeddings transfer cross-species function annotations to rice, maize, soybean, and Medicago more accurately than the autoencoder baseline. Subcellular localization, by contrast, remains dominated by sequence embeddings, consistent with the biology of the task, where targeting peptides and local motifs are encoded in sequence rather than coexpression context. We release the alignment pipeline and pre-aligned embeddings for the 5 seed species, providing a ready-to-use resource for plant functional genomics.

## Author Contributions

M.M. conceived the project and supervised the work. P.W. designed and implemented the Procrustes alignment pipeline, performed all experiments, analyzed the data, and wrote the first draft of the manuscript. J.M. provided the network and sequence data and helped improve the manuscript. All authors discussed the results and approved the final version.

## Acknowledgments

We thank the Mutwil lab for providing valuable comments on the project.

## Data and Code Availability

The pre-aligned embeddings for the 5 seed species and ORBIT code are publicly available at https://github.com/pwissenberg/orbit. UniProtKB/Swiss-Prot annotation counts were retrieved in April 2026 via the EBI Proteins API.

## Supplementary figures

**Figure S1.**
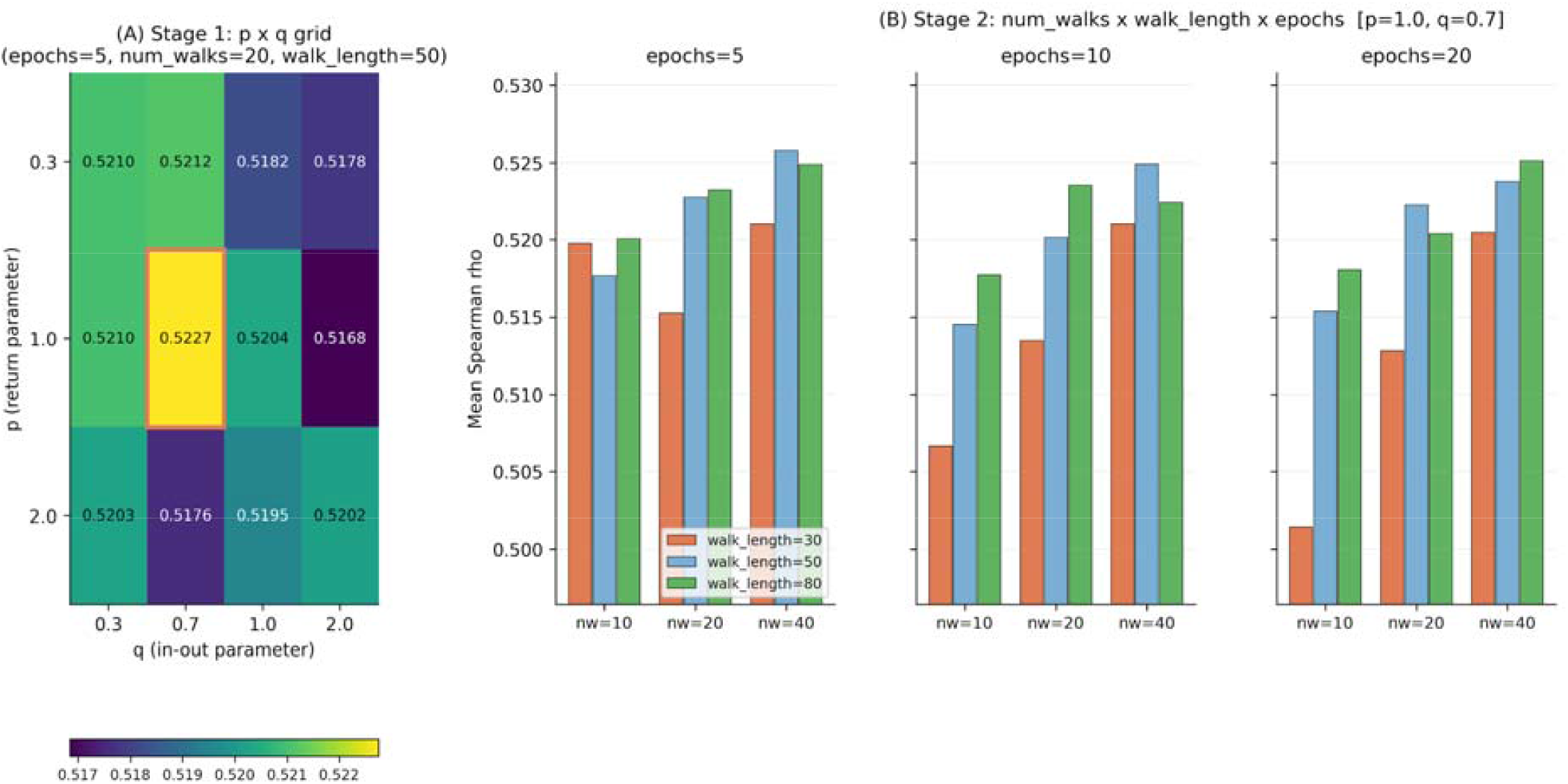
Node2Vec hyperparameter sensitivity. (A) Stage-1 grid search over return parameter p and in-out parameter q across 12 configurations, evaluated by cross-species Spearman correlation on three seed species pairs (ARATH-ORYSA, ARATH-BRADI, ORYSA-BRADI); the optimal combination (p = 1.0, q = 0.7) is highlighted. (B) Stage-2 grid search over number of walks, walk length, and training epochs across 27 configurations with p and q fixed at the Stage-1 optimum. Spearman ρ ranges from 0.502 to 0.526, indicating that embedding quality is robust to walk-structure parameters once p and q are tuned.

**Figure S2.**
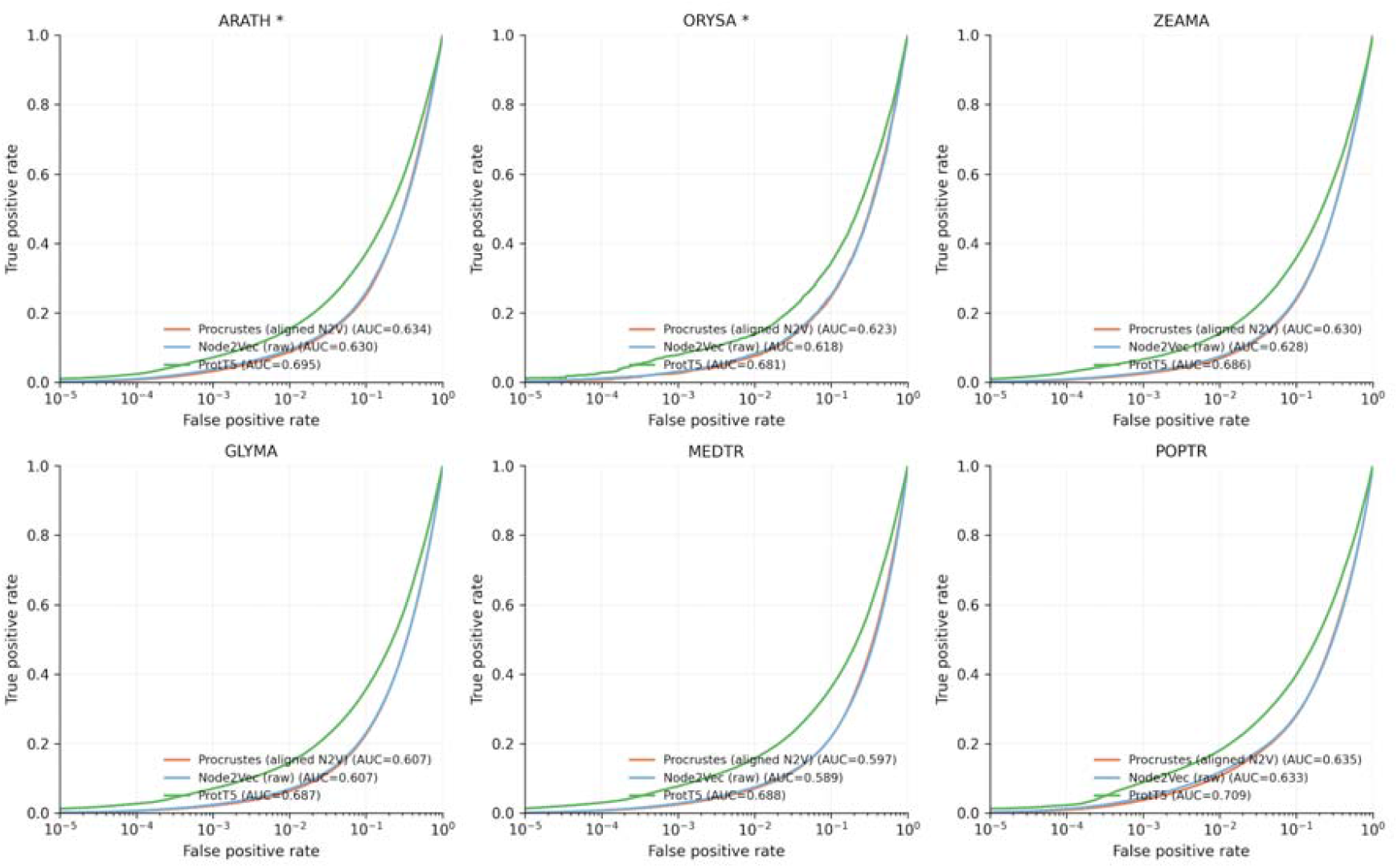
KEGG pathway co-membership ROC curves. Cumulative true-positives vs cumulative false-positives for gene pairs ranked by cosine similarity in the aligned embedding space, with KEGG pathway co-membership as ground truth, mirroring the SPACE evaluation protocol (Hu et al. 2025). Procrustes-aligned embeddings (orange) closely track the unaligned Node2Vec baseline (gray), confirming that alignment does not degrade pathway-level signal. ProtT5 sequence embeddings (green) outperform on every species (mean AUC ≈ 0.69 vs ≈ 0.62 for Procrustes), consistent with KEGG pathways being dominated by sequence-conserved enzyme families that ProtT5 captures directly. Of the five seed species, only ARATH and ORYSA have KEGG organism-code annotations available; PICAB, SELMO, and MARPO are not represented in KEGG and could not be evaluated. Asterisks denote seed species.

**Figure S3.**
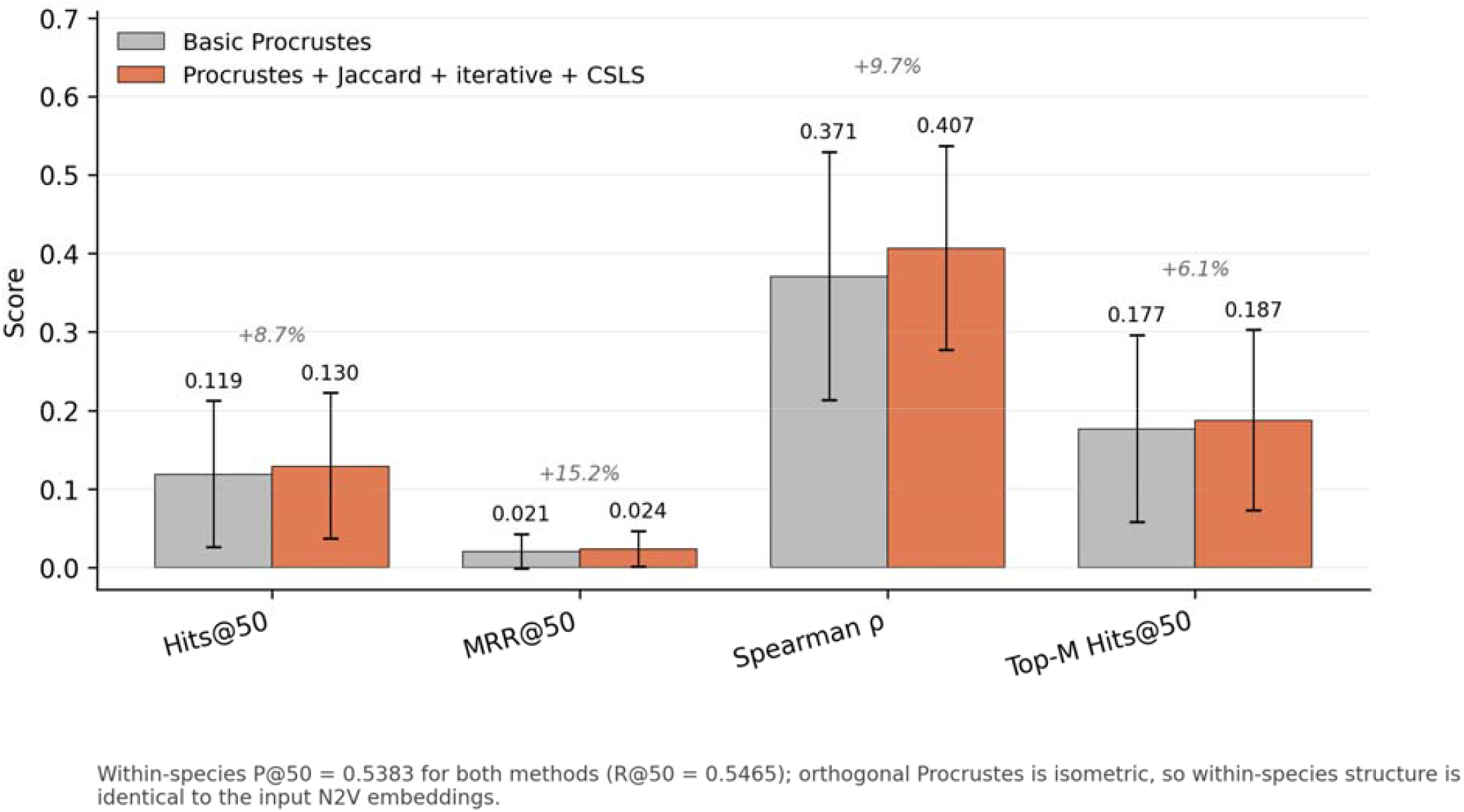
Component ablation of the Procrustes alignment pipeline. Comparison of basic orthogonal Procrustes against the full method (Procrustes + Jaccard-weighted ortholog pairs + iterative refinement + CSLS retrieval), both run on the same optimized Node2Vec embeddings (p = 1.0, q = 0.7). Cross-species metrics (Hits@50, MRR@50, Spearman ρ, Top-M Hits@50) isolate the contribution of the algorithmic improvements; within-species precision and recall are identical between the two variants by isometry, as expected for orthogonal rotations.

**Figure S4.**
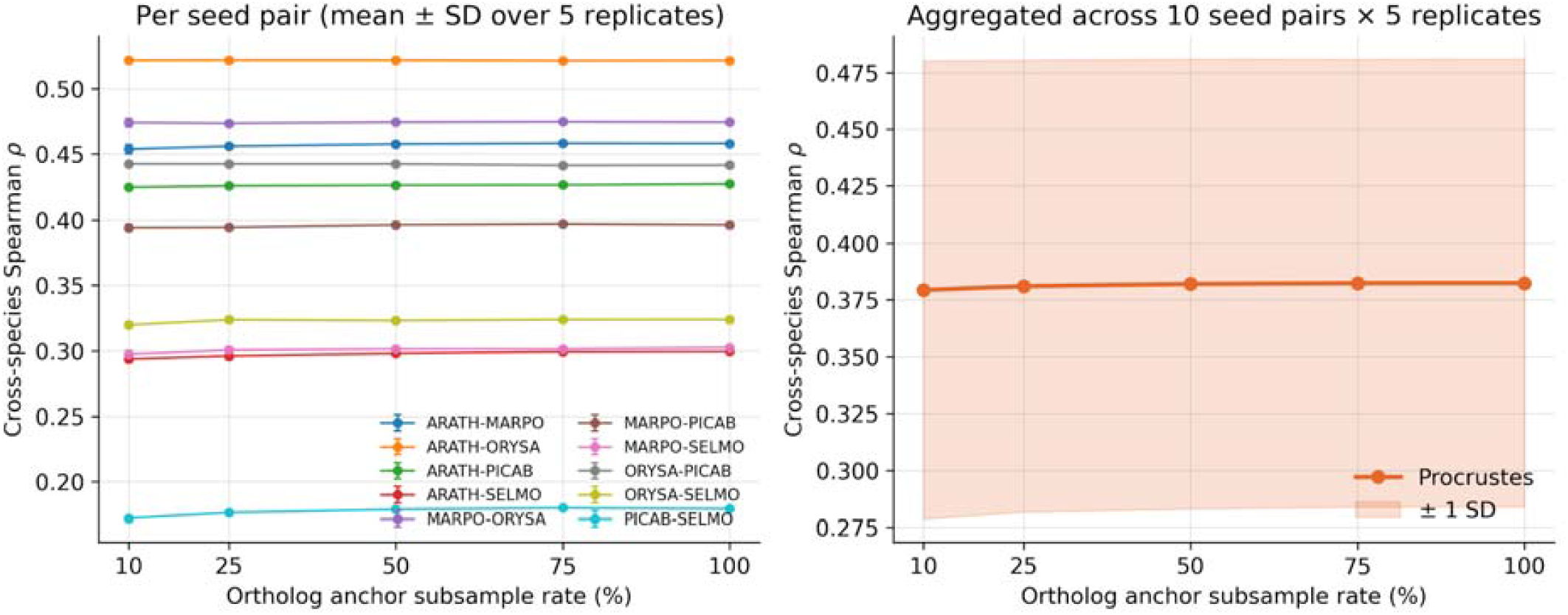
Robustness of Procrustes alignment to ortholog anchor subsampling. Cross-species Spearman correlation across all 10 seed-species pairs (ARATH, ORYSA, PICAB, SELMO, MARPO) as ortholog anchor pairs are randomly subsampled at rates of 100%, 75%, 50%, 25%, and 10%. Per-pair lines (light) and aggregated mean ± 1 SD shaded band (orange) across 5 random subsamples per rate. Mean Spearman ρ drops only 1.1% relative when anchors are reduced to 10% (0.382 → 0.379), and the largest single-pair drop (PICAB-SELMO, −4.0%) remains within one standard deviation across replicates, demonstrating that Procrustes alignment quality is essentially insensitive to ortholog count above the dimensional minimum.

**Figure S5.**
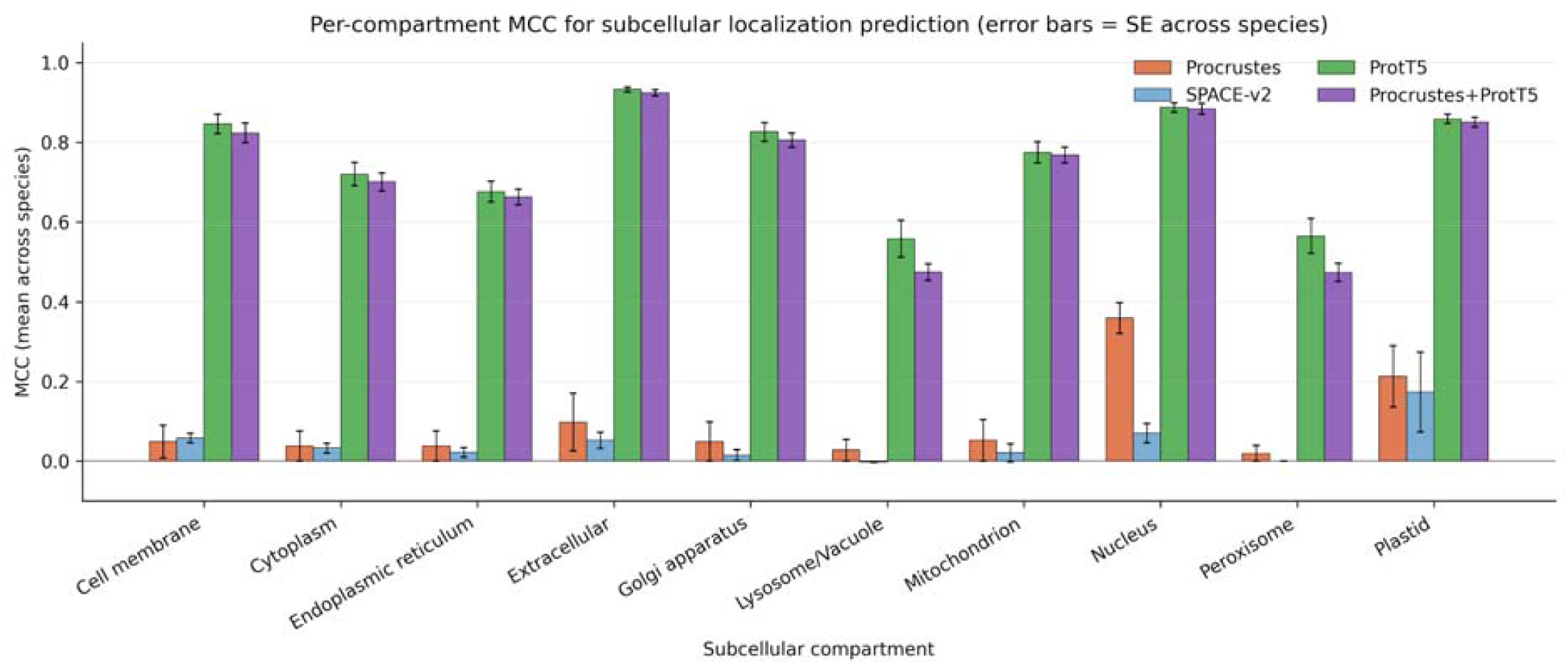
Per-localization Matthews correlation coefficient (MCC) on the DeepLoc 2.0 benchmark for Arabidopsis proteins. ProtT5 sequence embeddings (green) dominate localization classes encoded by N-terminal targeting peptides (Plastid, Mitochondrion, Secretory pathway, ER). Procrustes-aligned coexpression embeddings (orange) and SPACE-v2 (blue) perform comparably on Nucleus and Cytoplasm. Concatenating coexpression with sequence embeddings yields no improvement over ProtT5 alone for any compartment, consistent with the dominance of sequence-encoded targeting signals.

**Figure S6.**
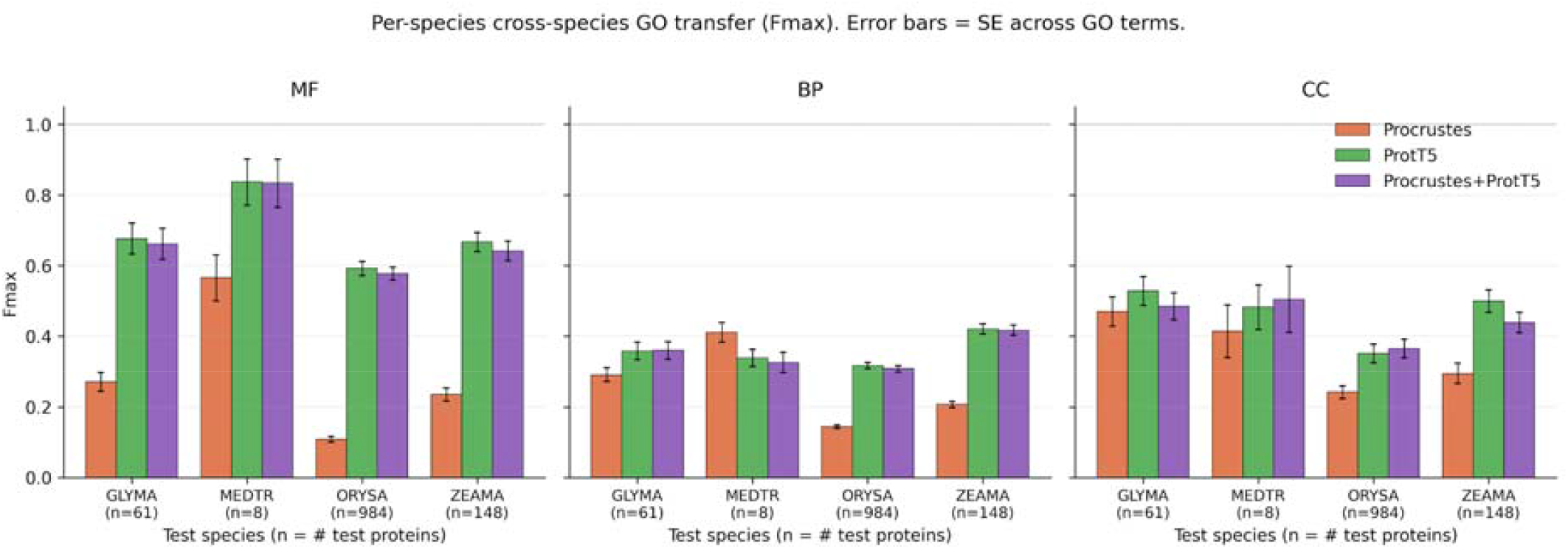
Per-species F-max for cross-species GO term transfer. Classifiers trained on A. thaliana experimental-evidence GO annotations (UniProt GOA) and tested on Oryza sativa, Zea mays, Glycine max, and Medicago truncatula. Results broken down by GO aspect (MF, BP, CC) and method (Procrustes, ProtT5, Procrustes + ProtT5 concatenation). Test-set sizes vary substantially across species (ORYSA n = 984, ZEAMA n = 148, GLYMA n = 61, MEDTR n = 8) and are annotated under each x-tick label. Error bars are standard errors computed across GO terms within each species/aspect combination 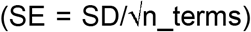, clipped to the valid F-max range [0, 1]; this treats terms as independent samples of task difficulty within an aspect.

## Supplementary tables

**Table S1. Node2Vec hyperparameter grid search**. Mean held-out edge AUC across {ARATH, ORYSA, PICAB} for 107 hyperparameter configurations spanning p ∈ {0.25, 0.5, 1, 2, 4}, q ∈ {0.25, 0.5, 0.7, 1, 2, 4}, num_walks ∈ {10, 20, 40}, walk_length = 50, epochs ∈ {3, 5, 10}. The (p, q) winner (p = 1.0, q = 0.7) was carried into Stage 2. Mean Spearman ρ between aligned embedding distance and orthogroup co-membership across pairs {ARATH–ORYSA, ARATH– BRADI, ORYSA–BRADI}, evaluated for 40 configurations at the Stage 1 winner (p = 1.0, q = 0.7), sweeping num_walks ∈ {10, 20, 40}, walk_length ∈ {30, 50, 80}, epochs ∈ {3, 5, 10}. The selected configuration (num_walks = 20, walk_length = 50, epochs = 10) was used for all reported experiments.

**Table S2. Full cross-species retrieval metrics per species pair**. Hits@50, MRR@50, Top-M Hits@50, and Spearman ρ for all 158 species pairs (10 seed pairs and 148 seed-to-non-seed pairs) under Procrustes (Jaccard-weighted + iterative + CSLS) and SPACE-v2 (autoencoder). Procrustes outperforms SPACE-v2 on every pair.

**Table S3. Runtime benchmarks for ORBIT and SPACE across 153 plant species**. Wall-clock time (in seconds) for each alignment method, decomposed into stage 1 (per-species setup, including Node2Vec embedding generation and any method-specific preprocessing) and stage 2 (pairwise cross-species alignment). total_seconds is the sum of stages 1 and 2. n_species is the number of species processed; n_success is the number of successful runs. Columns G–K report the distribution of per-species wall-clock time across all 153 species: mean, median, minimum, maximum, and standard deviation (in seconds). All benchmarks were run on a single CPU core under matched conditions.

**Table S4. DeepLoc 2.0 subcellular localization metrics per method**. Mean ± SD of F1-micro, F1-macro, accuracy, and Jaccard score across folds. CV = within-species 5-fold cross-validation; LOSO = leave-one-species-out across {ARATH, GLYMA, ORYSA, ZEAMA}.

**Table S5. NetGO 2.0 GO term prediction metrics per method**. F-max and AUPRC for Molecular Function (MF, n = 6 terms), Biological Process (BP, n = 21 terms), and Cellular Component (CC, n = 12 terms), evaluated on QuickGO experimental-evidence annotations.

