## Supplementary material for "ORBIT: Orthogonal Rotation for Biological Inter-species Transfer": Figure S1+6

Figure S1 — Node2Vec hyperparameter sensitivity (mean Spearman rho across ARATH-ORYSA, ARATH-BRADI, ORYSA-BRADI)

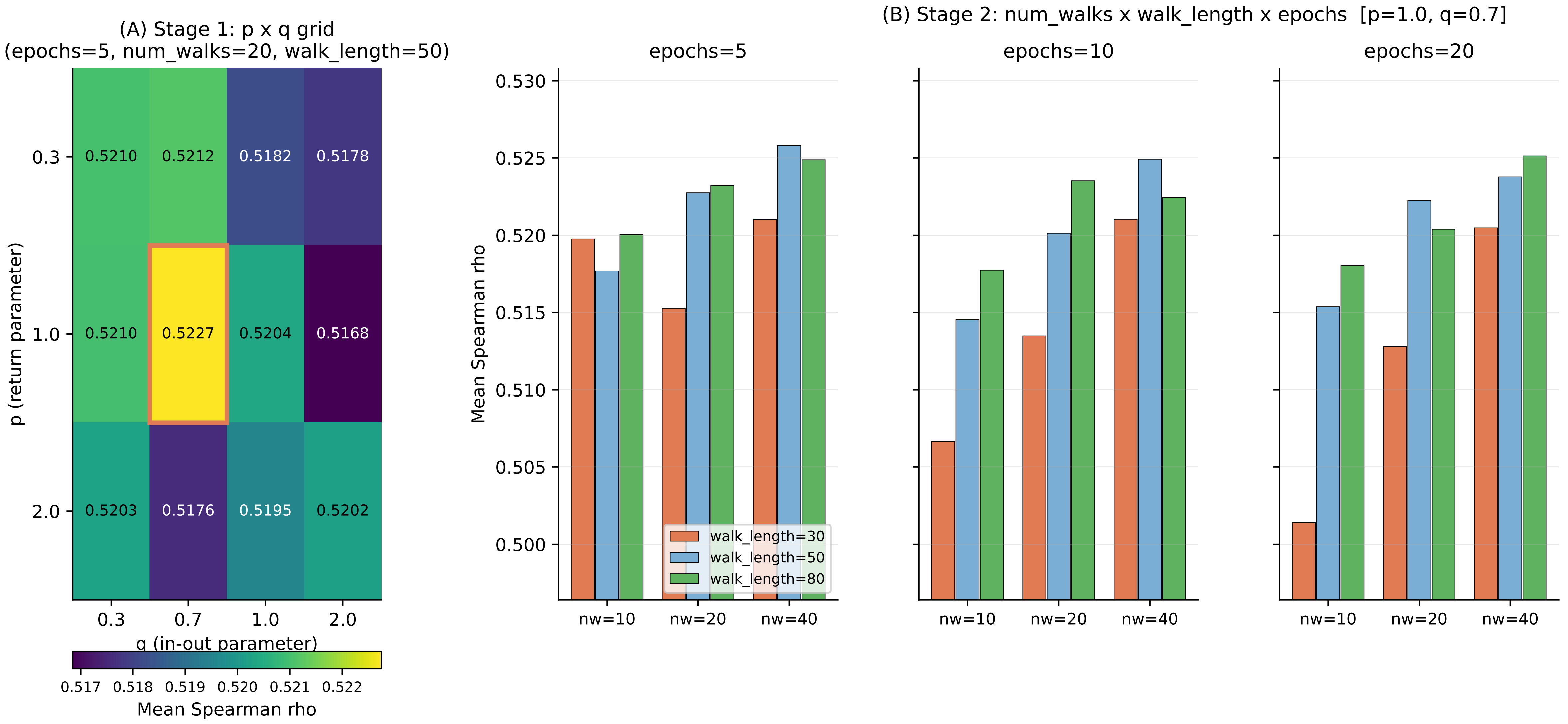

Figure S2 — KEGG pathway ROC curves (TPR vs FPR, log x). Y-axis normalized to TP / max_TP per species; legend AUCs are un-normalized full-AUC values. Asterisk (*) marks seed species.

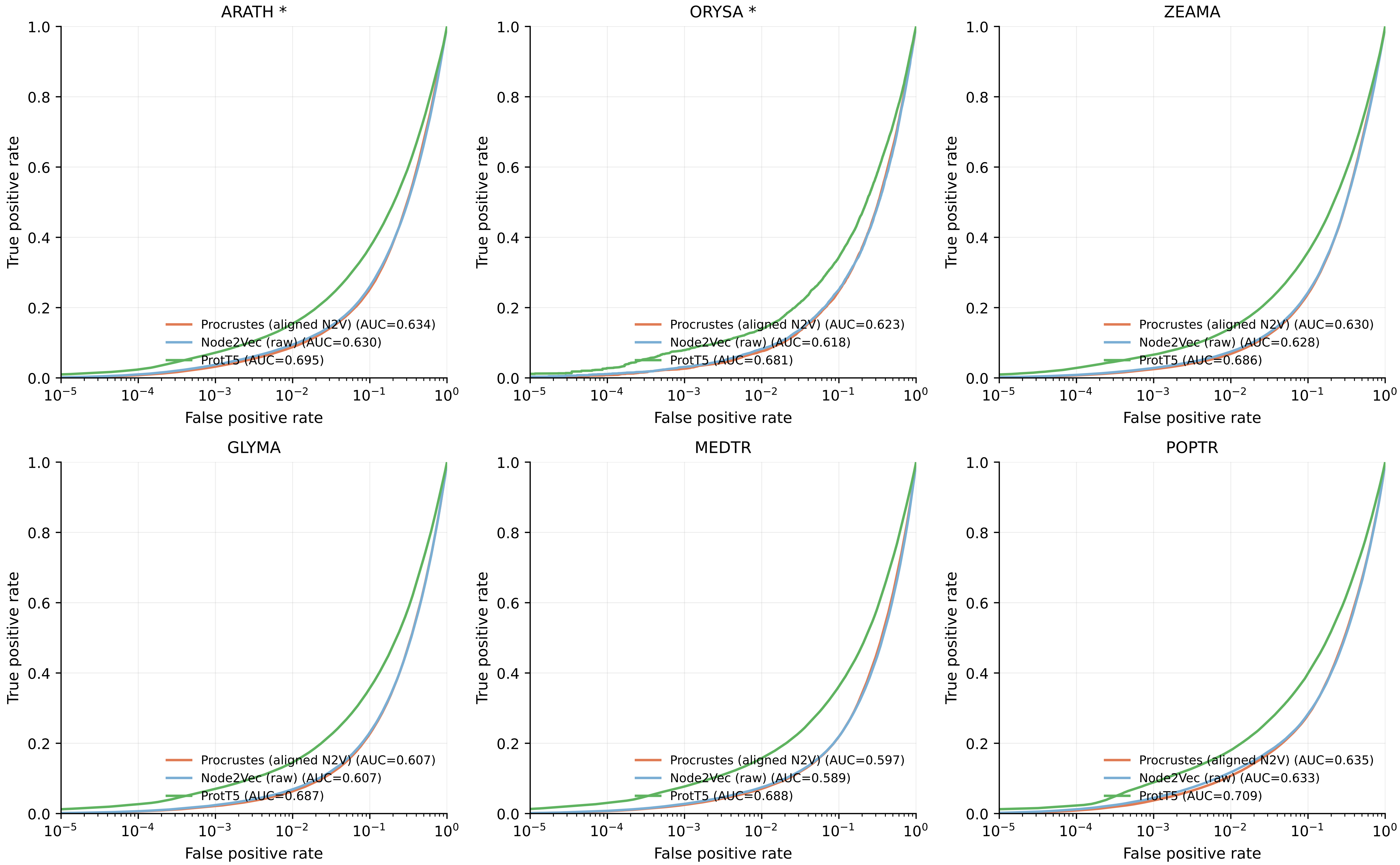

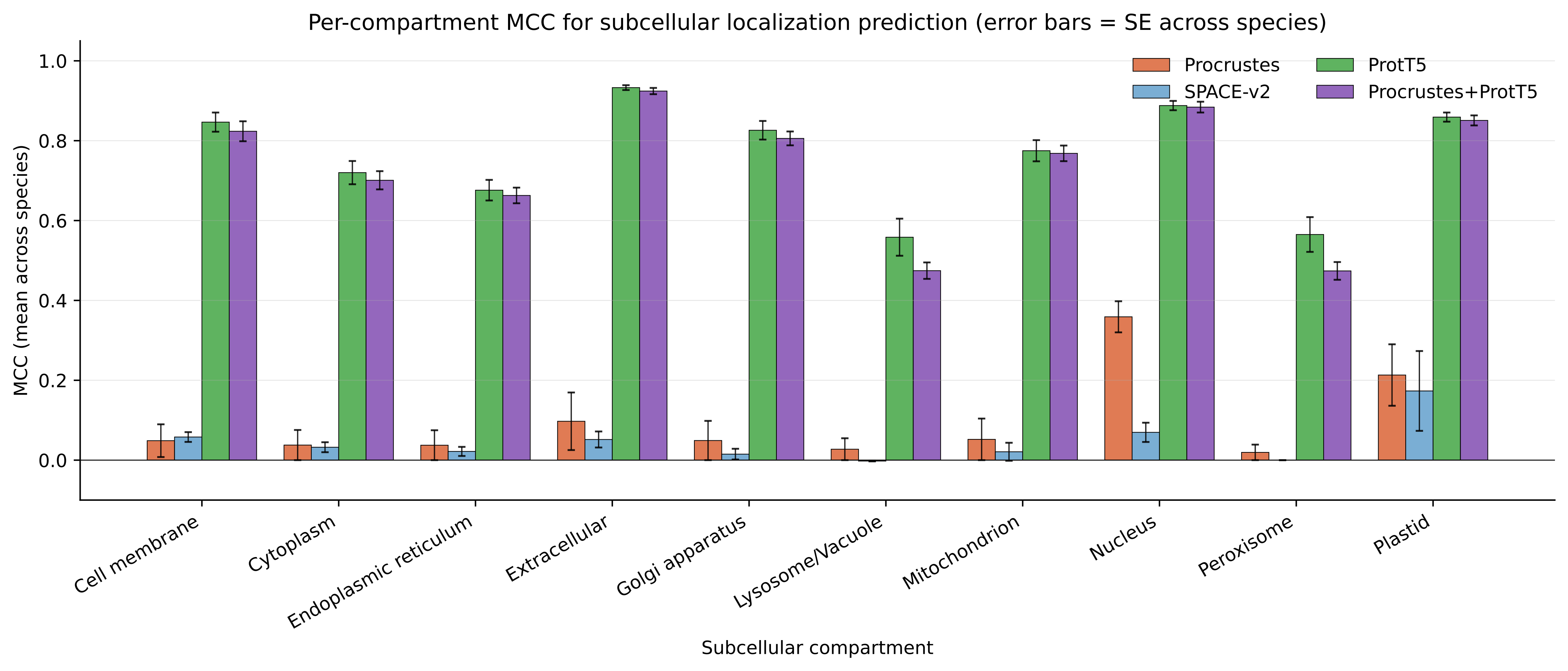
Per-compartment MCC for subcellular localization prediction (error bars = SE across species)
MCC (mean across species)
Subcellular compartment
Procrustes
SPACE-v2
ProtT5
Procrustes+ProtT5
Cell membrane
Cytoplasm
Endoplasmic reticulum
Extracellular
Golgi apparatus
Lysosome/Vacuole
Mitochondrion
Nucleus
Peroxisome
Plastid
0.0
0.2
0.4
0.6
0.8
1.0

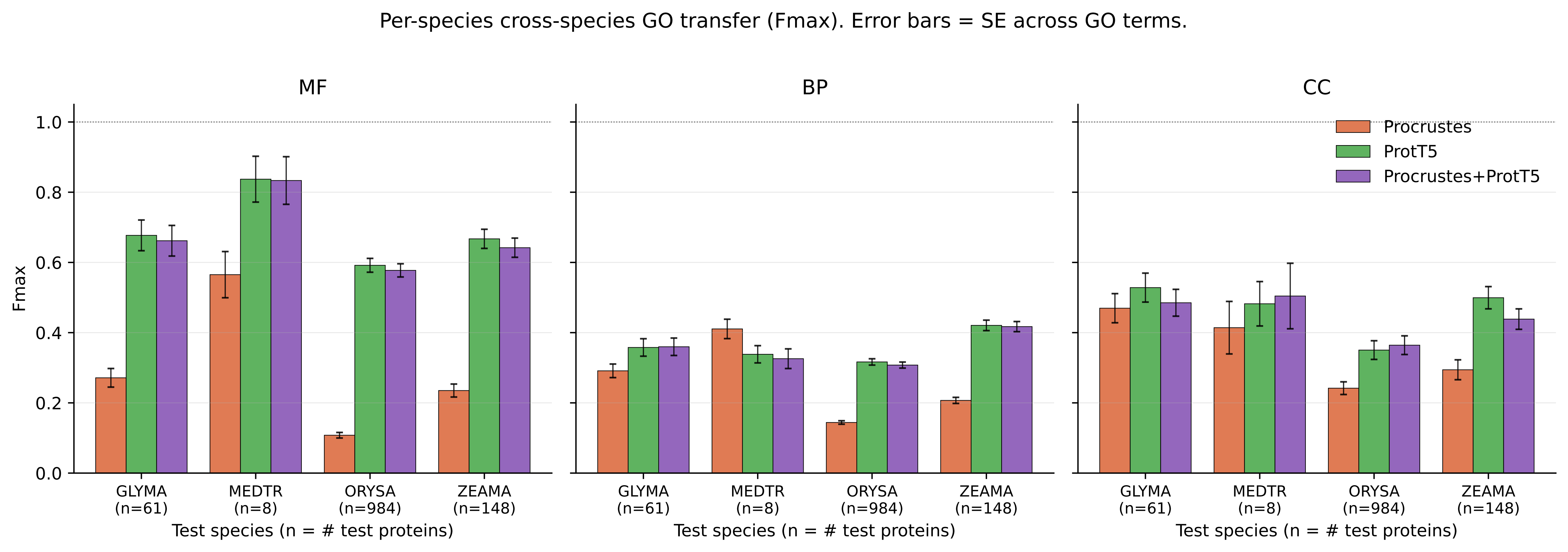
Per-species cross-species GO transfer (Fmax). Error bars = SE across GO terms.
MF
BP
CC
Fmax
1.0
0.8
0.6
0.4
0.2
0.0
Procrustes
ProtT5
Procrustes+ProtT5
GLYMA (n=61)
MEDTR (n=8)
ORYSA (n=984)
ZEAMA (n=148)
Test species (n = # test proteins)

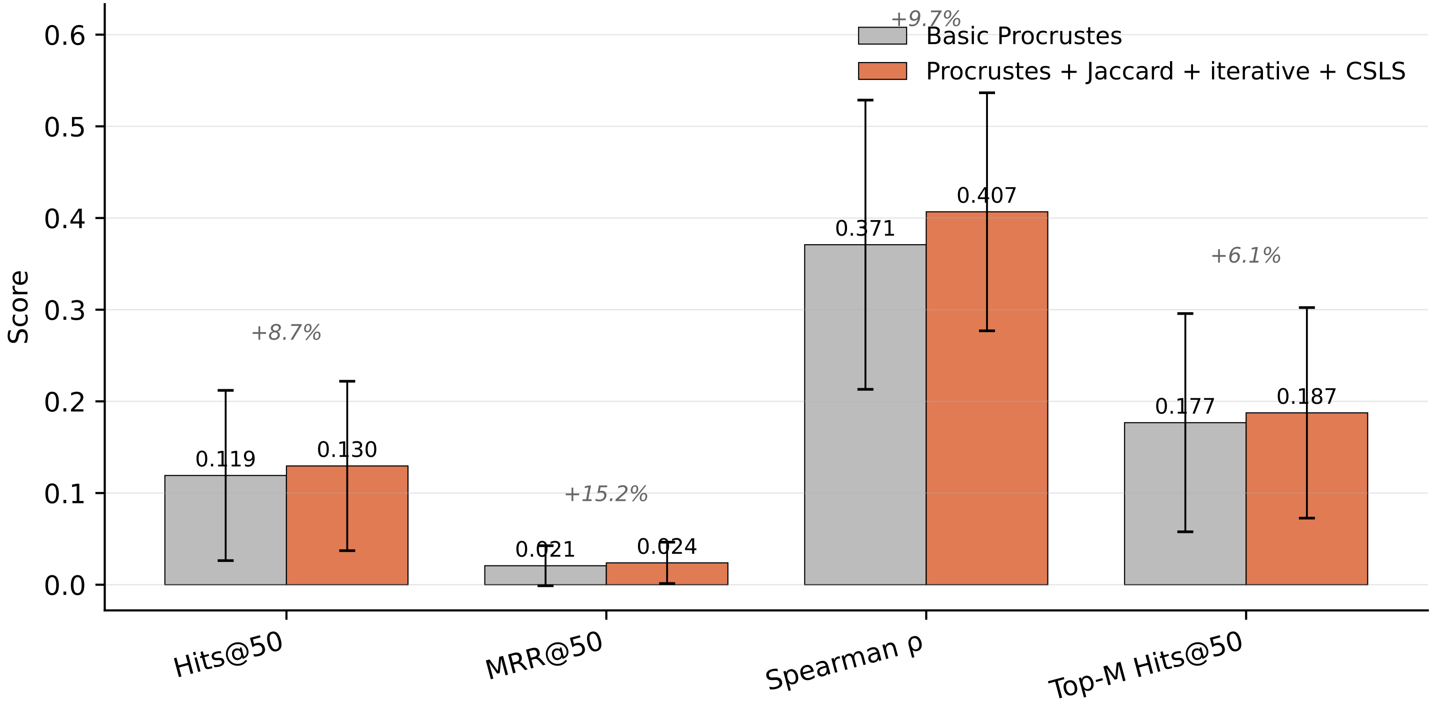


Within-species P@50 = 0.5383 for both methods (R@50 = 0.5465); orthogonal Procrustes is isometric, so within-species structure is identical to the input N2V embeddings.

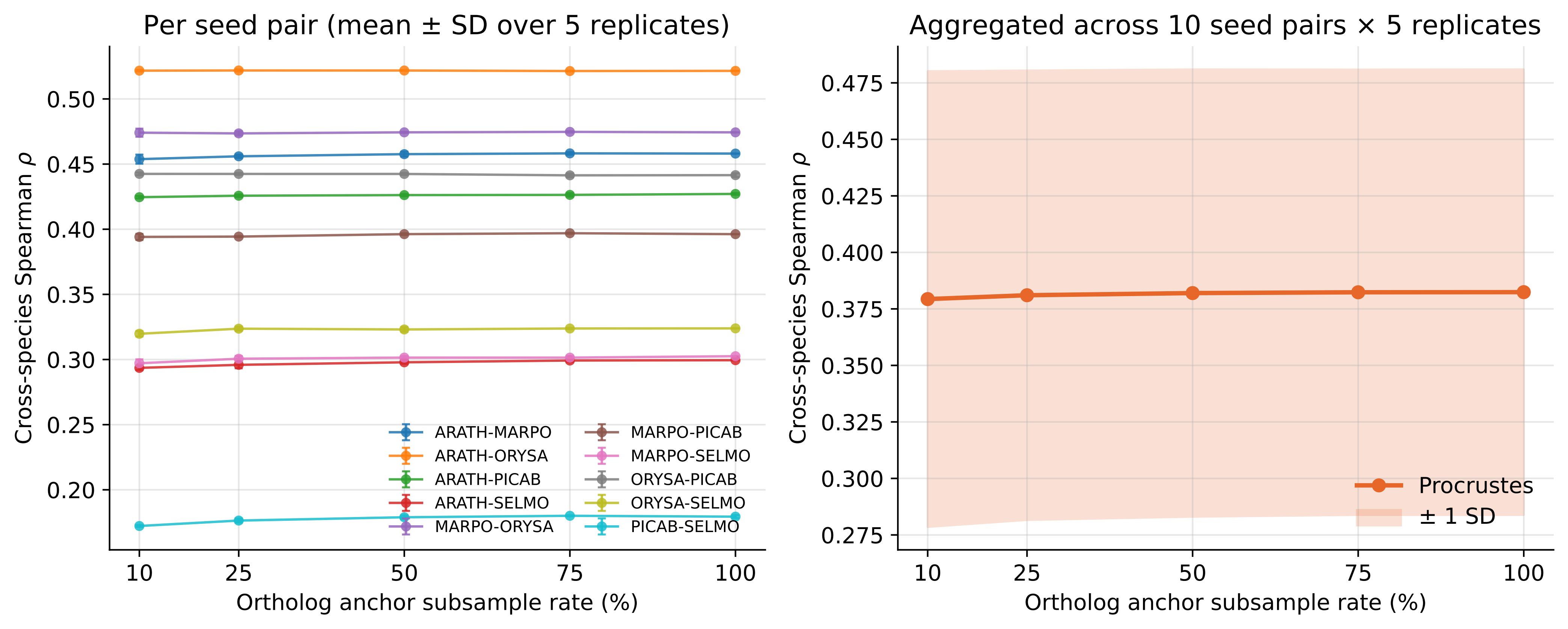
Per seed pair (mean ± SD over 5 replicates)
Aggregated across 10 seed pairs × 5 replicates
Cross-species Spearman ρ
Ortholog anchor subsample rate (%)
ARATH-MARPO
ARATH-ORYSA
ARATH-PICAB
ARATH-SELMO
MARPO-ORYSA
MARPO-PICAB
MARPO-SELMO
ORYSA-PICAB
ORYSA-SELMO
PICAB-SELMO
Procrustes
± 1 SD
